# Tissue geometry encodes a surface tension gradient that drives epithelial renewal

**DOI:** 10.64898/2026.04.28.721444

**Authors:** Zhang Wen, Michael P. Murrell, Kaelyn Sumigray

## Abstract

The mammalian intestinal epithelium renews itself every few days through coordinated cell movement and extrusion along the crypt-villus axis, yet how these processes are physically organized across intact tissue remains unknown. Using long-term live imaging of intact intestinal tissue, nuclear strain mapping, and targeted perturbations of cell-cell adhesion, actomyosin contractility, and extracellular matrix adhesion, we find that epithelial cells experience a graded mechanical landscape encoded by villus geometry; tension is highest at the villus tip and decays toward the base. This tissue-scale tension gradient, generated by E-cadherin-mediated adhesion and actomyosin contractility, is the dominant predictor of collective cell velocity, outperforming force magnitude and cell density. Artificial re-establishment of a tension gradient is sufficient to reorient epithelial migration, whereas disruption of the gradient arrests both cell movement and extrusion. Epithelial renewal further requires a precise balance between this tension gradient and extracellular matrix-derived friction; excessive friction uncouples migration from extrusion, leading to pathological cell accumulation. Together, these findings reveal that intestinal epithelial homeostasis is organized by a geometry-encoded mechanical gradient that coordinates collective cell motion, extrusion, and tissue repair through a single tissue-scale framework.

---

The intestinal epithelium undergoes rapid and continuous renewal, with cells born in the crypt migrating collectively along the villus axis before being eliminated at the tip (*1–4*). Despite decades of study, the physical mechanisms that coordinate this large-scale epithelial migration across intact tissue remain unclear. At the tissue scale, this process resembles a steady epithelial flow, persisting despite constant cell turnover (*5–7*). In particular, it is unclear what physical mechanism organizes collective migration over hundreds of microns (*8–10*) while maintaining tight coupling to epithelial cell loss and tissue integrity during homeostasis (*4*, *6*, *11–13*).

Live imaging studies have established that villus epithelial cells migrate actively rather than passively, and that cells near the villus tip move faster than those at the base (*14*). These observations support models in which position-dependent active migration supplements mitotic pressure from crypts to drive epithelial flow. However, such frameworks do not explain how epithelial motion is coordinated across the entire villus, nor how migration remains robust in the face of continuous extrusion and mechanical perturbation. More broadly, it remains unresolved whether large-scale epithelial flow is governed primarily by cell-intrinsic migratory programs or instead emerges from tissue-scale physical constraints.

The intestinal villus exhibits a highly stereotyped and conserved three-dimensional architecture, characterized by a narrow, highly curved tip that gradually transitions to a broader base (*11*, *13*, *25*). In other 3D epithelial systems, geometry is mechanically coupled to surface tension and stress (*9*, *11*, *16*, *19–22*), suggesting that villus architecture itself could encode a long-range mechanical gradient. We therefore hypothesized that epithelial migration along the villus is organized by a geometry-encoded gradient in mechanical tension (*9*, *13*, *16*, *26–28*).

In many biological systems, coherent tissue movements can arise from spatial gradients in surface tension rather than from local propulsion alone, generating long-range flows analogous to Marangoni flows (*15–18*). Such flows arise when geometry and material properties together generate a driving force that biases motion along the surface tension gradient (*9*, *11*, *16*, *19–22*). Notably, epithelial tissues are capable of sustaining surface tension through the combined action of cell-cell adhesion and actomyosin contractility, raising the possibility that similar principles could operate during tissue renewal (*23*, *24*).

In physical systems governed by Marangoni flow, material motion does not depend on absolute force magnitude but instead emerges from a balance between spatial gradients in surface tension and resistance to motion. A defining feature of such systems is that velocity (*v*) scales with the surface tension gradient (∇*γ*) and is attenuated by friction (*ξ*), allowing the driving and dissipative contributions to be tested independently (*v*∼∇*γ*⁄*ξ*). If epithelial migration along the villus is governed by such a mechanism, then epithelial velocity should scale with the magnitude of the tissue-scale tension gradient and be suppressed by increased extracellular resistance.

Here, we show that villus geometry generates a robust tissue-scale surface tension gradient, highest at the tip and decaying toward the base, which can be read out by nuclear strain (*ε*) within epithelial cells. This mechanically patterned landscape depends on E-cadherin-mediated adhesion and actomyosin contractility and predicts both the magnitude and direction of collective epithelial motion. These data are consistent with Marangoni-like flows regulating cell movement up the villus, where velocity is set by the balance between a surface tension gradient and friction (*16*, *22*). Perturbing the gradient collapses coordinated flow, while enhancing it amplifies tissue-scale motion and accelerates epithelial reorganization following cell loss. We further show that epithelial flow and renewal depend not only on this gradient, but also on an optimal level of extracellular matrix-derived friction, which constrains motion and extrusion. Together, our results reveal that intestinal epithelial renewal is encoded by tissue architecture, rather than arising solely from local or cell-intrinsic migratory programs.

## Intestinal epithelial cells exhibit coherent, directional flows along the villus axis

To directly visualize epithelial motion across intact villi, we performed long-term, high-resolution live imaging of mouse intestinal villi using an ex vivo tissue slice culture system that preserves native crypt-villus architecture. This approach allowed continuous imaging of epithelial dynamics over extended periods (Fig 1A-A’) while maintaining physiological cell turnover (Fig 1B-C; Movies S1-S3) and tissue integrity. Tracking epithelial cells revealed coherent, directional motion toward the villus tip (Movie S4). Particle image velocimetry (PIV) analysis confirmed that trajectories were highly aligned over tens to hundreds of microns along the villus axis (Fig 1D), indicating that epithelial migration occurs as a coordinated tissue-scale flow rather than as independent or locally confined movements.

**Figure 1.**
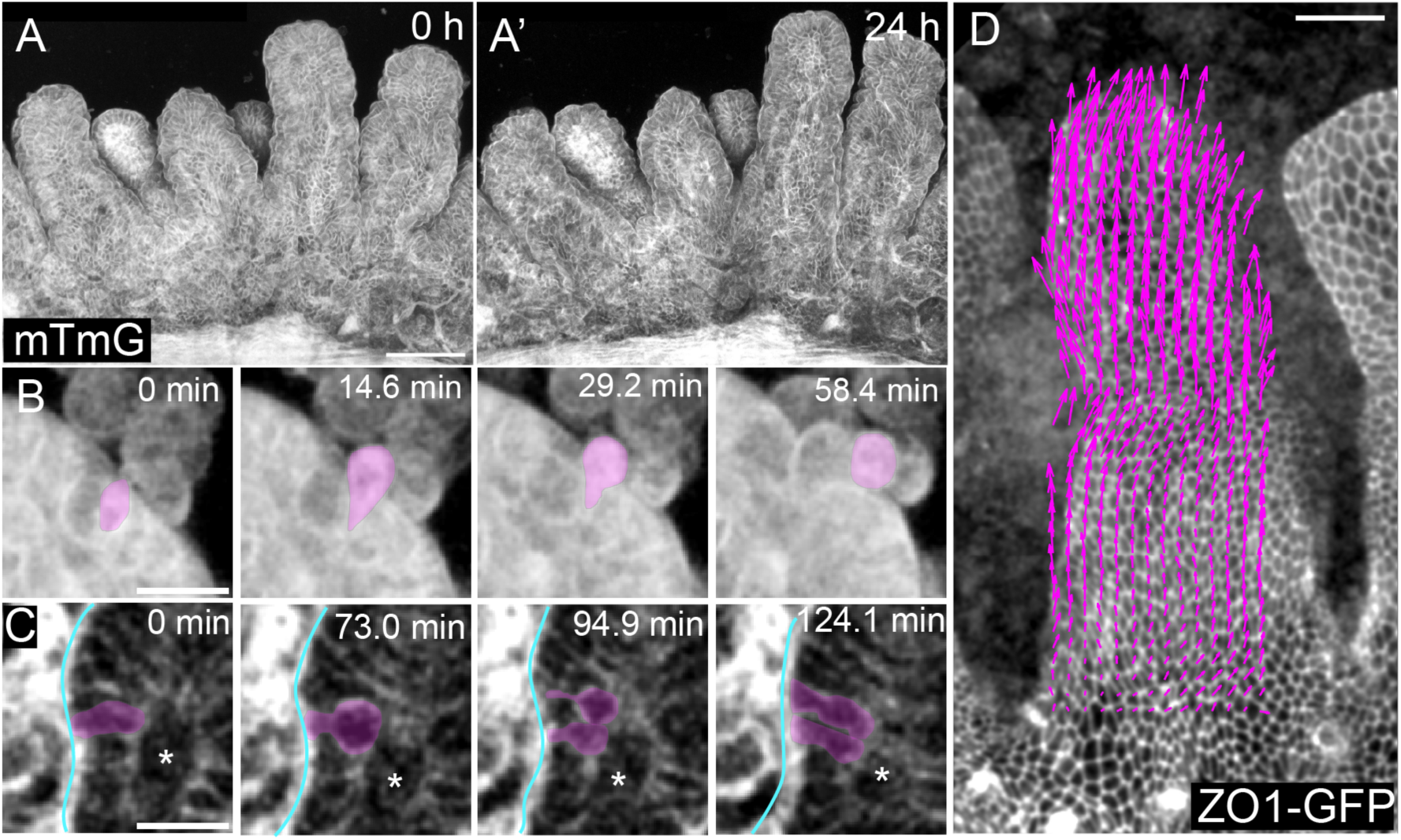
Long-term live imaging reveals directional epithelial flow in the villus. **A)** Representative confocal images of intact intestinal slice from mTmG mice imaged from 0 h to 24 h of live tissue culture, demonstrating preservation of villus architecture and epithelial integrity over extended imaging. Scale bar, 200 µm. **B-C)** Representative confocal images of time-lapse live imaging showing cell extrusion at the villus tip and cell proliferation in the crypt. Scale bars, 20 µm. **D)** Long-term live imaging of intact intestinal tissue reveals sustained, directional epithelial migration along the villus axis. Scale bar, 50 µm.

To quantify the spatial organization of this epithelial flow, we measured cell velocity as a function of position along the villus axis. Consistent with previous reports (*14*), epithelial velocity progressively increased as cells moved toward the villus tip (Fig 2A-C). Continuous imaging revealed that this position-dependent velocity profile was stable over time and maintained across villi despite ongoing cellular turnover, indicating that epithelial flow is not uniform but instead exhibits a robust spatial gradient along the villus.

**Figure 2.**
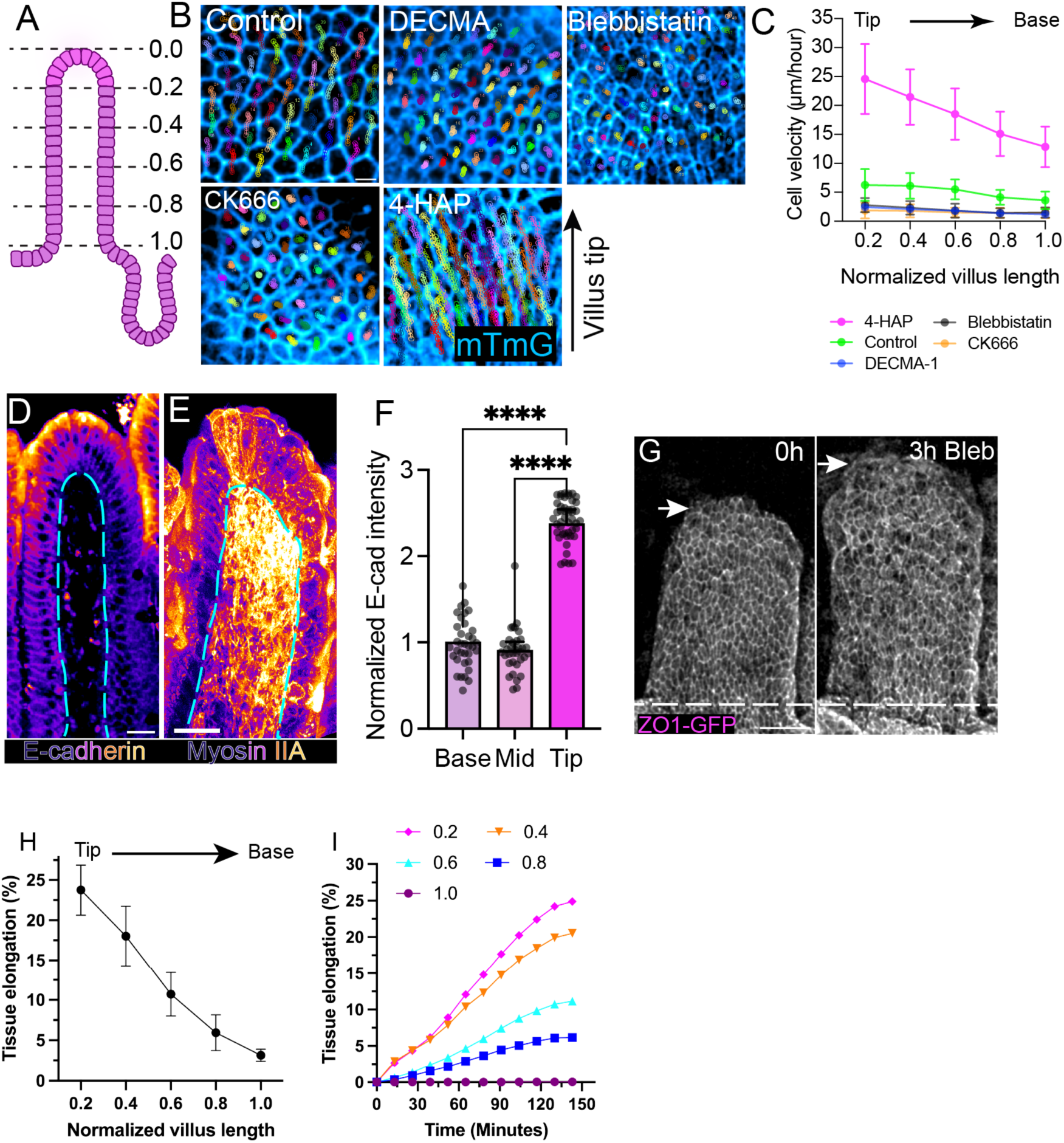
Cell-cell adhesion and actomyosin contractility sustain directional epithelial flow in the villus. **A)** schematic indicating normalized positional bins along the villus axis used for velocity quantification, with 0 representing the villus tip and 1 the villus base. **B)** Representative cell trajectories over 3 h under control conditions and following perturbation of adhesion (DECMA-1), contractility (blebbistatin), branched actin assembly (CK666), or enhanced myosin II activity (4-HAP). Colors denote individual tracked cells. Scale bar, 10 µm. **C)** Mean epithelial cell velocity as a function of normalized villus position under indicated conditions (n=9 to 10 villi from N=3 mice). **D-E)** Representative heat maps of E-cadherin (D) and non-muscle myosin IIA (E) localization and intensity along the villus epithelium. Dashed line marks the basement membrane. Scale bars, 20 µm. **F)** Quantification of normalized E-cadherin intensity at the base, mid-villus, and tip (n=35 villi from N=3 mice). **G)** Representative images of tissue elongation upon blebbistatin over 3 hours. Scale bar, 50 µm. **H)** Total tissue elongation as a function of normalized villus position. ***I*)** Tissue elongation as a function of time for each villus position. Data are mean ± SD. Multigroup comparisons used one-way analysis of variance with a Tukey test.

## Epithelial flow velocity is regulated by a tissue-scale tension gradient

Surface tension in epithelia arises from the combined action of cell-cell adhesion and actomyosin contractility (*16*, *22*, *23*, *30*, *31*). We therefore asked whether these force-generating components are spatially patterned along the villus axis in a manner consistent with a tissue-scale mechanical gradient. Whole-mount imaging revealed enrichment of both E-cadherin and non-muscle myosin IIA at the villus tip (Fig 2D-F). Their graded distribution aligned with the direction and increasing velocity of epithelial flow.

Because cell-cell adhesion and actomyosin contractility are the principal determinants of epithelial surface tension, these graded patterns suggested the presence of a tissue-scale tension gradient along the villus axis (*16*, *23*). To test whether adhesion- and contractility-dependent tension is required to sustain epithelial flow, we perturbed these components while monitoring collective cell motion for several hours in intact tissue. Disrupting E-cadherin-mediated adhesion (DECMA-1), inhibiting myosin II contractility (blebbistatin), or blocking Arp2/3-dependent actin assembly/active migration (CK666) did not simply reduce velocity uniformly; instead, each perturbation abolished the spatial organization of epithelial motion and stalled flow across the villus (Fig 2B-C; Movies S5-S7). In contrast, enhancing myosin II activity with 4-HAP (*32*, *33*) increased epithelial cell velocity and amplified directional flow toward the villus tip (Fig 2B-C; Movie S8). Consistent with a gradient of pre-existing tension, acute inhibition of myosin II caused rapid, position-dependent tissue relaxation, with villus tips elongating more than basal regions (Fig 2G-I; Movie S9). Together, these data demonstrate that epithelial motion along the villus occurs as a coherent, spatially graded flow that depends on intact cell-cell adhesion and contractility. We next asked whether this organization reflects an underlying tissue-scale mechanical gradient capable of coordinating epithelial motion.

## Villus geometry generates the tissue-scale surface tension gradient

The intestinal villus exhibits a stereotyped geometry that is conserved across regions of the small intestine: a highly curved tip that transitions to a broader, flatter base (Figure 3A). Because curvature is mechanically coupled to surface tension in diverse biological systems (*9*, *11*, *19–21*, *34*), we hypothesized that villus architecture itself generates a long-range mechanical gradient capable of organizing epithelial behavior. To test this, we first quantified villus shape and confirmed that curvature is highest at the villus tip and decreases along the sides (Figure 3B).

**Figure 3.**
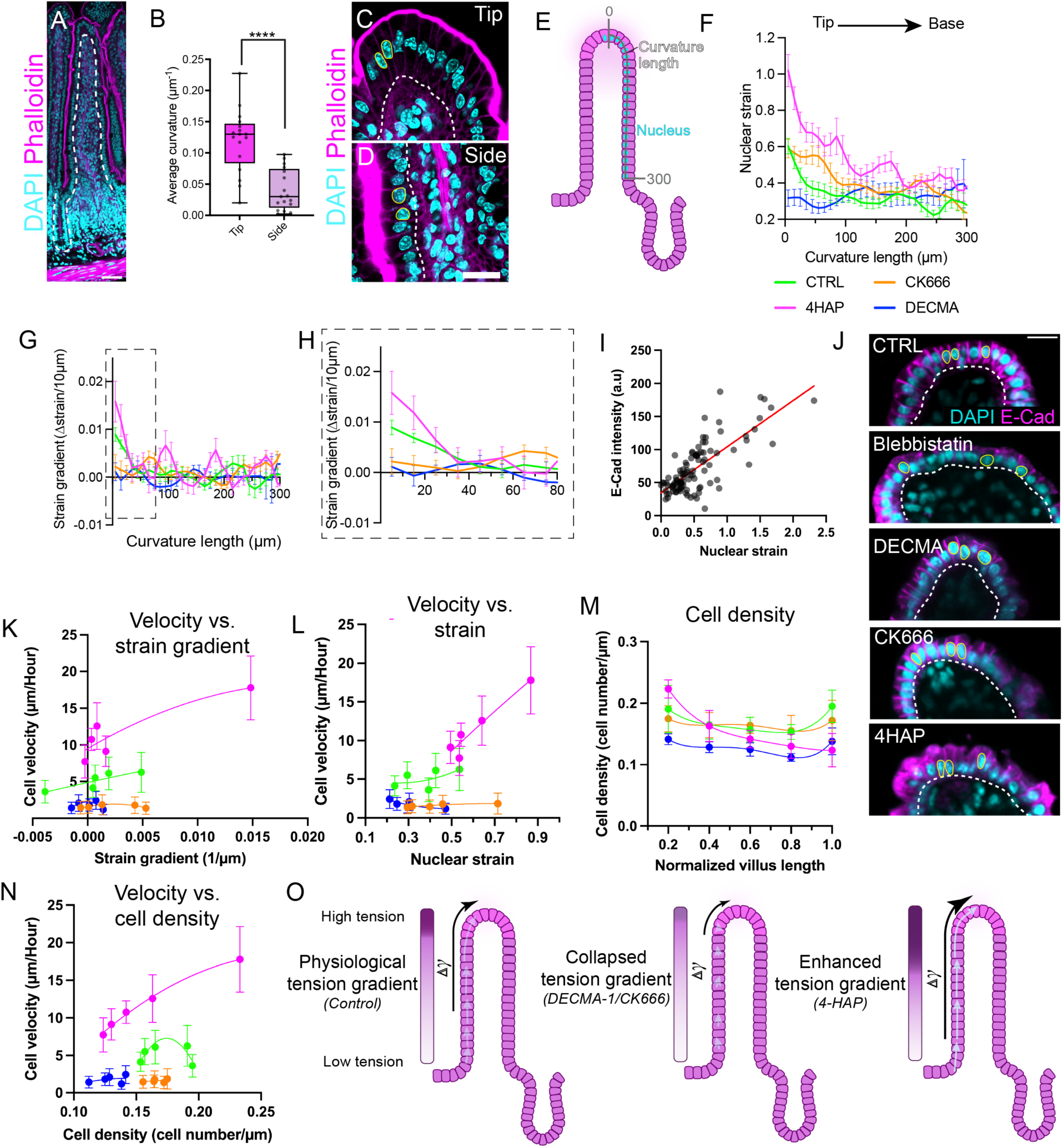
A tissue-scale tension gradient drives directional epithelial flow in the villus. **A)** Whole-mount confocal image of an adult mouse small intestine (jejunum) stained for F-actin (phalloidin, magenta) and nuclei (DAPI, gray), illustrating a villus defining the tip region analyzed throughout the study. **B)** Quantification of average epithelial curvature at the villus tip vs. lateral side regions (n=17 villi from N=3 mice). **C, D)** Representative confocal images of villus epithelium stained for F-actin (magenta) and nuclei (DAPI, cyan). White dashed line marks basement membrane. Scale bars, 20 µm. **E)** Schematic illustrating measurement of nuclear strain along the curvature length of the villus epithelium, with distance measured from the villus tip (0 µm). **F)** Nuclear strain plotted as function of curvature length for each condition. Control (CTRL, green) revealing a graded decay from the villus tip toward the base in control. (n=15 villi from N=3 mice for each condition). Data are mean ± SEM. **G)** Strain gradient, measured as the change in strain over every 10 µm, plotted against curvature length, revealing a steep gradient at the villus tip (n=17 villi from N=3 for each condition). Data are mean ± SEM. **H)** Expanded view of strain gradients near the villus tip (boxed region in G), highlighting modulation of gradient magnitude by adhesion and contractility. **I)** Correlation between E-cadherin intensity and nuclear strain. Red line indicates linear fit. R^2^ = 0.5732, P<0.0001. **J)** Representative villus tip stained for E-cadherin (magenta) and nuclei (cyan) under control conditions and following perturbation of cell-cell adhesion (DECMA-1), contractility (blebbistatin), branched actin assembly (CK666) or enhanced myosin II activity (4-HAP). White line marks basement membrane. Scale bar, 20 µm. **K)** Cell velocity plotted against local strain gradient across all conditions, revealing a strong positive association between collective motion and tension gradient magnitude. **L)** Cell velocity plotted as a function of nuclear strain. M) Cell density as a function of normalized villus position under indicated conditions. N) Cell velocity plotted against local cell density. **O)** Summary schematic illustrating how collapse, maintenance, or enhancement of the tissue-scale tension gradient corresponds to disrupted, physiological, or accelerated collective epithelial migration, respectively. Across all perturbations, epithelial velocity correlates with the local strain gradient rather than with absolute nuclear strain or cell density, consistent with Marangoni-like flow in which motion is driven by spatial gradients in surface tension rather than force magnitude.

We next asked whether this geometric gradient corresponds to a mechanical stress gradient within the epithelium. Nuclear shape is a sensitive reporter of intracellular force, with nuclear elongation reflecting increased mechanical strain (*35–38*). Nuclear strain gradients therefore provide a proxy for spatial differences in epithelial tension. Mapping nuclear morphology along the villus axis revealed that nuclei at the tip were markedly more elongated than those at the base (Fig 3C-D). When plotted as a function of absolute position along the curved villus axis (Fig 3E), nuclear strain (ε = AR − 1) decayed smoothly from tip to base (Fig 3F), establishing the presence of a large-scale tissue strain gradient (Δ*ε*⁄Δ*x*) spanning the villus (Fig 3G-H).

To determine whether this strain gradient reflects spatial patterning of epithelial surface tension, we examined the molecular contributors to intercellular force generation. E-cadherin fluorescence intensity, which is enriched at the villus tip, positively correlated with nuclear strain (Fig 3I), suggesting that cell-cell adhesion contributes to the observed mechanical landscape. Together, these data demonstrate that villus geometry is translated into a graded mechanical field across the epithelium, providing a physical substrate capable of biasing collective cell behavior over long distances.

To test whether this tissue-scale strain is actively generated, we examined how it responds to perturbations of force-generating machinery. Inhibition of E-cadherin-mediated adhesion (DECMA-1) or branched actin assembly (CK666) abolished the spatial decay of nuclear strain along the villus, collapsing the gradient (Fig 3G-H, J; Fig S1A-B). Conversely, enhancing myosin II activity with 4-HAP increased both the magnitude and spatial extent of the strain gradient, extending the characteristic length scale over which elevated strain was maintained (Fig 3G-H; Fig S1A-B).

We next asked which mechanical parameter best predicts epithelial velocity. Across all conditions, cell velocity correlated most strongly with the strain gradient, and only moderately with absolute nuclear strain (Fig 3K-L, Table 1). By contrast, cell density did not correlate with velocity (Fig 3M-N, Table 1), indicating that, unlike models in which density profiles reflect the outcome of active migration, cell velocity here is more directly predicted by the strain gradient (*14*). Notably, in 4-HAP-treated tissues, where both contractility and the strain gradient are amplified, velocity strongly correlated with multiple mechanical parameters. In contrast, in DECMA- and CK666-treated tissues, where the strain gradient collapses, these correlations were lost (Fig 3K-L, Table 1). Together, these analyses indicate that the spatial gradient of tension, rather than force magnitude alone, is the dominant predictor of collective epithelial motion (Fig 3O).

**Table 1.** Correlation analysis between cell velocity and nuclear strain, strain gradient, cell density.

| R<sup>2</sup> value | CTRL | 4-HAP | DECMA-1 | CK666 |
| --- | --- | --- | --- | --- |
| Nuclear strain | 0.3131 | 0.9199 | 0.7200 | 0.6000 |
| Strain gradient | 0.7544 | 0.8018 | 0.0003 | 0.0006 |
| Cell density | 0.0060 | 0.9761 | 0.1347 | 0.1457 |

To test whether a tension gradient is not only necessary but sufficient to orient collective migration, we locally restored myosin II activity in otherwise contractility-suppressed tissue using regionalized photo-inactivation of blebbistatin (*39*, *40*). Reestablishing a spatial gradient of contractility was sufficient to rapidly reorient epithelial motion toward the region of higher tension, even in the absence of endogenous patterning (Fig 4A-C). This locally induced gradient generated coherent directional flow comparable to control tissues, as confirmed by PIV and velocity measurements.

**Fig. 4.**
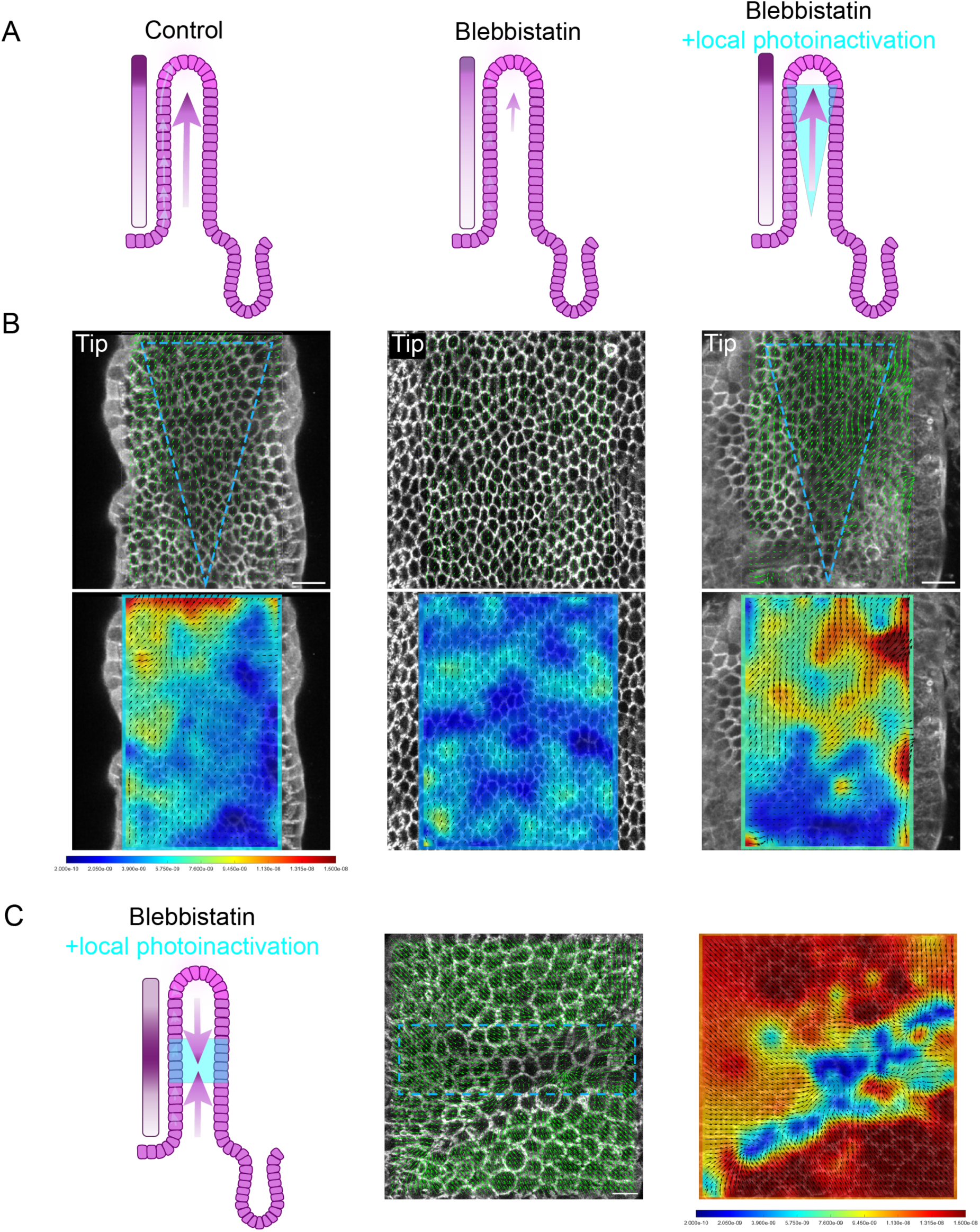
Local restoration of contractility reestablishes a tension gradient and reorients epithelial flow. **A)** Schematic illustrating the experimental design: live tissues were globally treated with blebbistatin, followed by local blue light illumination to photo-deactivate blebbistatin and restore contractility. The illuminated region of interest (ROI, triangular shape) reestablished a local tension gradient comparable to control and blebbistatin treated conditions. **B)** Representative PIV analyses of cell motion in control and blebbistatin-treated tissues. After photo-deactivation, collective cell motion reappeared, recapitulating the gradient observed in control. Green arrows, velocity vector. Heatmap, velocity magnitude. Scale bars, 20 µm. **C)** PIV analysis showing that cells surrounding the ROI moved directionally toward the photo-deactivated region following restoration of contractility. Scale bar, 20 µm.

## Extracellular matrix-mediated friction gates tension-driven epithelial flow

In classical Marangoni systems, flow velocity is determined not only by the magnitude of the surface tension gradient but also by resistance to motion (*29*). We therefore asked whether ECM-mediated friction modulates how the villus tension gradient is interpreted by migrating epithelial cells. To selectively increase friction without directly altering contractility, we enhanced integrin-ECM adhesion using manganese, which stabilizes integrin binding (*41–44*). We tested two levels of increased friction: a moderate condition (6 µM Mn^2+^; M6) and a high-friction condition (20 µM Mn^2+^; M20). Increasing ECM friction markedly reduced epithelial flow velocity in a dose-dependent manner, despite preservation of the villus tension gradient (Fig 5A-B). Cell velocities were reduced across the villus axis under both M6 and M20 conditions, with the strongest suppression observed at the villus tip (Fig 5B, Movies S10-S11). Notably, nuclear strain profiles remained elevated at the villus tip under increased friction (Fig 5C-D), indicating that tissue tension was not lost. Notably, while the strain gradient remained intact in moderate friction conditions, it collapsed under high friction conditions (Fig 5E-F), further supporting that cell motion is driven by the strain gradient and not absolute strain.

**Figure 5.**
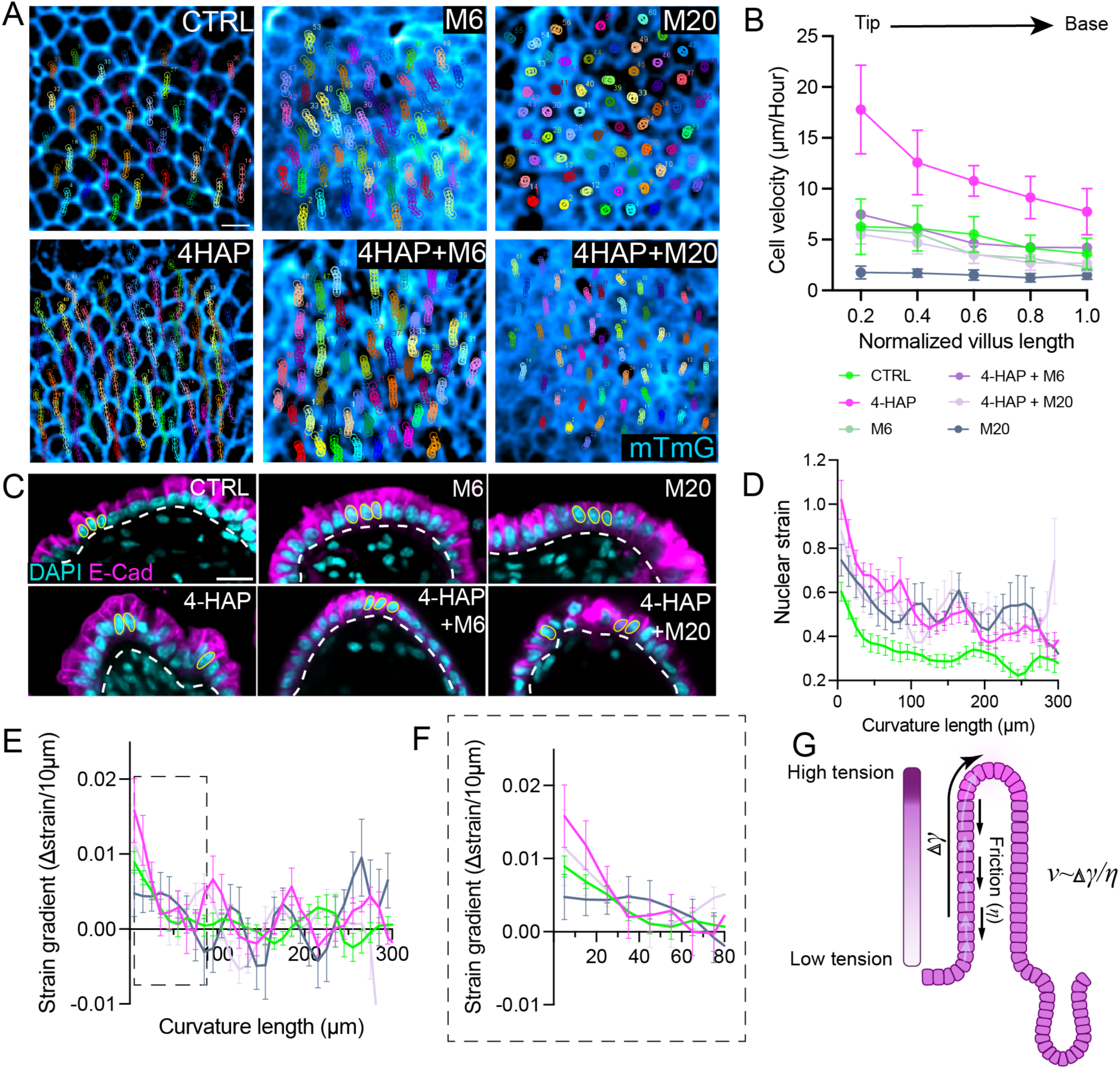
Excessive friction impairs epithelial flow in the villus. **A)** Representative cell trajectory overlays illustrating collective epithelial motion under indicated conditions. Scale bar, 10 µm. **B)** Mean epithelial cell velocity plotted as a function of normalized villus position along the crypt-villus axis, with 0 representing the villus tip and 1 the villus base. **C)** Representative high magnification images of villus tips showing nuclear morphology across conditions. Scale bar, 20 µm. **D)** Nuclear strain plotted as a function of curvature length under control, high-friction (M20), high-tension (4-HAP), and combined conditions (n=15 villi from N=3 mice). Data are mean ± SEM. **E)** Strain gradient plotted along the villus curvature for indicated conditions, showing attenuation of the gradient under high friction (n=15 villi from N=3 mice). Data are mean ± SEM. **F)** Expanded view of strain gradients near the villus tip, highlighting loss of gradient magnitude under excessive friction. **G)** A schematic illustrating the Marangoni force balance, in which epithelial velocity scales with the ratio of tissue-scale surface tension gradient to ECM-derived friction.

To test whether increased tension could compensate for elevated friction, we combined manganese treatment with myosin II activation (4-HAP). Enhancing contractility partially rescued epithelial migration under moderate-friction conditions (4-HAP+M6) but failed to restore flow under high-friction conditions (4-HAP+M20 ≈ M20 alone) (Fig 5A-B; Movies S12-S13). These results indicate that while increased tension can overcome moderate resistance, excessive friction fundamentally limits tissue-scale flow. Together, these findings demonstrate that epithelial motion along the villus is governed by a balance between a driving tension gradient and ECM-derived friction (Fig 5G). Tension alone is insufficient to sustain effective flow when resistance is high, revealing friction as a critical mechanical gate that modulates tension-driven epithelial dynamics. In systems governed by Marangoni-like flow, velocity reflects the balance between a driving tension gradient and resistive friction (*v*∼ ∇*γ*⁄*ξ*). Our perturbations map directly onto this minimal force balance. Manipulations that reduce the tissue-scale surface tension gradient, through disruption of E-cadherin adhesion, inhibition of actomyosin contractility, or suppression of actin-based force organization, abolish both the strain gradient and collective epithelial flow. Conversely, enhancing myosin II activity increases the magnitude and spatial extent of the gradient and proportionally amplifies epithelial velocity. In contrast, increasing integrin-mediated adhesion elevates friction without eliminating the tension gradient, resulting in a dose-dependent suppression of flow that can be partially overcome by increased contractility only under moderate resistance. These results demonstrate that epithelial velocity is controlled by the ratio of a geometry-encoded tension gradient to extracellular friction, consistent with Marangoni-like force balance rather than with models based on absolute force magnitude, cell density, or local migratory activity alone (Fig 5G). However, epithelial renewal requires not only directed migration, but also continuous and spatially coordinated cell extrusion at the villus tip. Therefore, we next asked how disrupting this mechanical balance impacts epithelial turnover and tissue homeostasis.

## A balanced tension-friction landscape enables rapid tissue reorganization during homeostasis

To determine how the tissue-scale tension gradient contributes to epithelial renewal, we quantified epithelial turnover directly by measuring cell extrusion events during long-term live imaging of intact villi (Fig. 6A-B, Movie S14). Perturbations that altered the tension gradient produced corresponding changes in epithelial extrusion. Enhancing actomyosin contractility with 4-HAP significantly increased extrusion frequency, whereas disrupting cell-cell adhesion (DECMA-1) or inhibiting branched actin assembly (CK666) markedly reduced extrusion relative to control tissues (Fig 6B). These effects mirrored changes in strain gradient magnitude and spatial extent observed under the same perturbations, suggesting that epithelial extrusion is mechanically coupled to the tissue-scale tension gradient.

**Figure 6.**
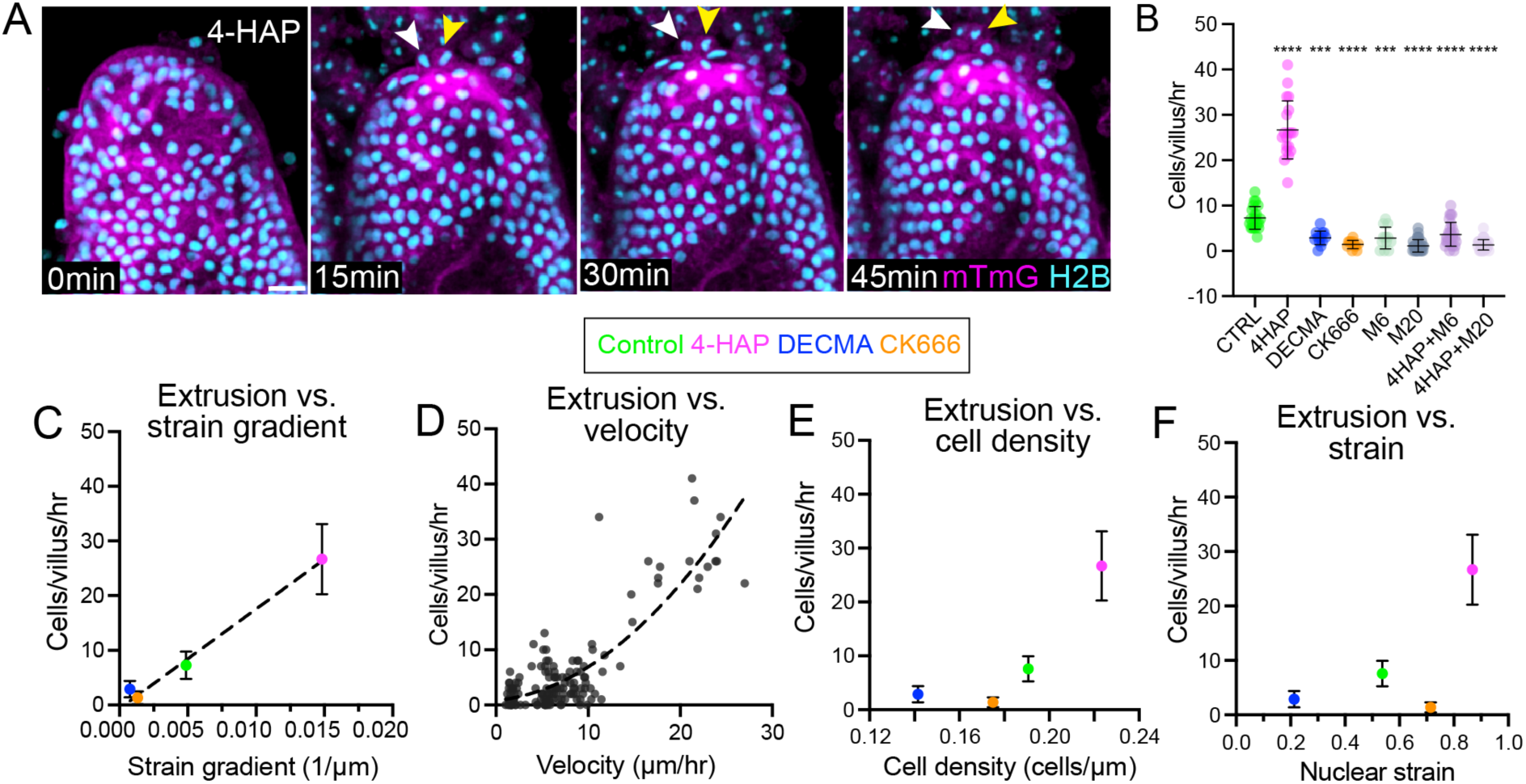
A balanced tension-friction landscape couples epithelial flow to extrusion during homeostasis. **A)** Time-lapse images of villus tips from mTmG; H2b-GFP-labeled tissue treated with 4-HAP, showing dynamic epithelial extrusion events over time. Arrowheads denote extruding cells. Scale bar, 20 µm. **B)** Quantification of epithelial extrusion rate (cells per villus per hour) under control conditions and following perturbation of contractility (4-HAP), cell-cell adhesion (DECMA-1), branched actin assembly (CK666), increased cell-ECM adhesion (manganese; M6 and M20), and combined manganese and myosin activation (4-HAP+M6, 4-HAP+M20). Data are mean ± SD (n=11 to 31 villi from N=3 mice for each condition). ****P<0.0001, ***P=0.0001. **C)** Mean extrusion rate plotted as a function of the local strain gradient, revealing a strong association between extrusion frequency and tension gradient magnitude. Extrusion rate correlates most strongly with the tissue-scale strain gradient, indicating that spatial differences in tension, rather than velocity, force magnitude, or density, determine where epithelial loss occurs in Marangoni-like tissue flow. **D)** Relationship between epithelial cell velocity and extrusion rate across all conditions. Dashed line indicates nonlinear fit. **E)** Mean extrusion rate plotted as a function of local cell density. **F)** Mean extrusion rate plotted as a function of nuclear strain under indicated conditions. Multigroup comparisons used one-way analysis of variance with a Tukey test.

To identify which mechanical parameter best predicts epithelial extrusion, we correlated extrusion rates with cell velocity, nuclear strain, and the local strain gradient at the villus tip, where extrusion predominantly occurs. Strikingly, extrusion rate correlated most strongly with the strain gradient (Fig. 6C-F, Table 2). Thus, epithelial extrusion is governed primarily by spatial differences in tension rather than by force magnitude or cell movement alone.

**Table 2.**
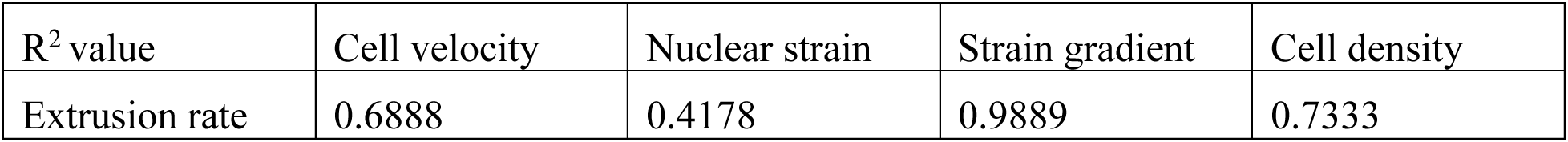
Correlation analysis between extrusion rate and nuclear strain, strain gradient, cell density.

Because epithelial renewal requires extrusion to be coordinated with upward cell motion, we next asked whether extracellular resistance alters this coupling. Under conditions of elevated ECM-mediated friction, extrusion rates were disproportionately suppressed relative to migration (Fig 6B), leading to progressive accumulation of cells along the villus (Fig S2). Increasing contractility with 4-HAP restored epithelial velocity under these conditions but failed to rescue extrusion, demonstrating that excessive friction uncouples migration from cell loss despite preserved tissue tension (Fig. 5B and Fig. 6B). These findings indicate that epithelial homeostasis depends on a finely tuned balance between a driving tension gradient and permissive friction.

To test whether the same tension-friction landscape governs epithelial reorganization following acute cell loss, we induced controlled, localized epithelial damage using tissue-scale laser ablation to mimic acute extrusion events (Fig. 7A, Movie S15). In control villi, ablation at the tip triggered rapid tissue recoil followed by progressive wound closure that resolved within ∼15 minutes, consistent with efficient redistribution of tensile forces and coordinated cell movement (Fig. 7B, Fig. S5). In tissues treated with 4-HAP, initial recoil was significantly larger and wound closure occurred more rapidly, accompanied by increased velocity of neighboring cells, consistent with elevated tissue tension enhancing the capacity for mechanical reorganization (Fig. 7C, Movie S16). By contrast, in DECMA-treated villi, where the tissue-scale tension gradient is disrupted, ablation produced limited recoil and wounds failed to close, leaving persistent epithelial gaps (Fig. 7A-C; Movie S17). Under these conditions, neighboring cell velocity was markedly reduced, demonstrating that impaired collective motion compromises tissue renewal.

**Fig. 7.**
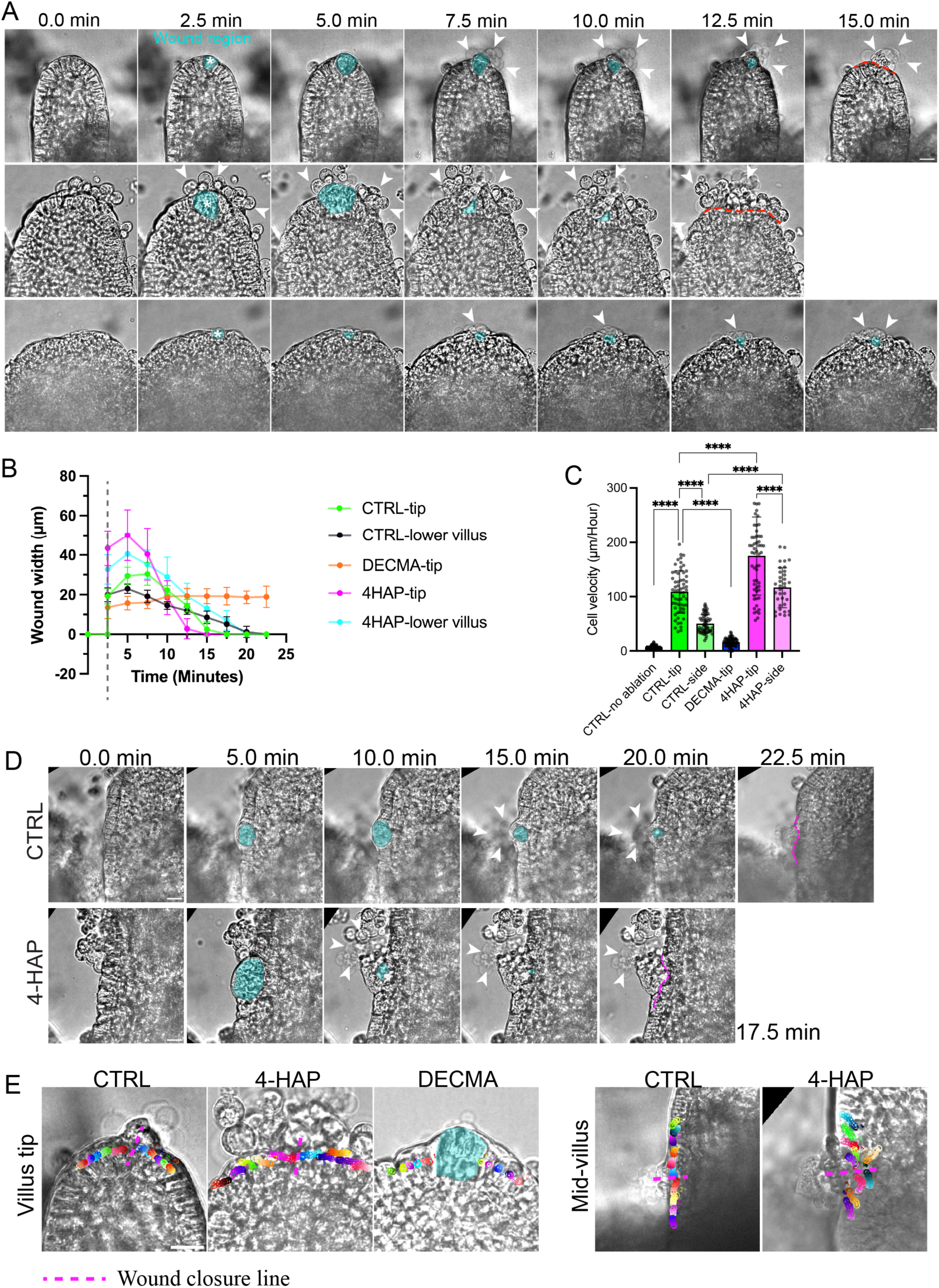
Surface tension gradients coordinate epithelial repair following cell loss. **A)** Time-lapse images following localized laser ablation (asterisk) at the villus tip. showing tissue recoil and wound closure (cyan) dynamics in control, 4-HAP-treated, and DECMA-treated tissues. Arrowheads indicate extruding cells, dashed magenta line marks closed wound. Scale bar, 20 µm. **B)** Quantification of wound width over time following laser ablation at the villus tip or along the villus side under indicated conditions. Dashed line marks time of laser ablation (n=7 villi from N=3 mice). **C)** Quantification of epithelial cell velocity adjacent to the wound region following laser ablation (n=7 villi from N=3 mice). ****P<0.0001. **D)** Representative time-lapse images showing laser-induced tissue damage at the mid-villus region and subsequent wound opening and closure under control and 4-HAP–treated conditions. Scale bar, 10 µm. **E)** Representative cell trajectories during wound closure following laser ablation. Scale bar, 10 µm. Data are mean ± SD. Multigroup comparisons used one-way analysis of variance with a Tukey test.

Consistent with spatial patterning of tension along the villus, ablation along the villus side resulted in slower wound closure and reduced cell velocity compared to ablation at the tip, although both were enhanced by 4-HAP treatment (Fig 7B-E). Together, these results demonstrate that the same Marangoni-like force balance between surface tension gradients and friction governs steady-state epithelial turnover and acute tissue repair, linking migration, extrusion and reorganization through a common mechanical framework.

## Discussion

The mammalian small intestinal epithelium renews itself through continuous collective cell migration and extrusion, yet how these processes are physically coordinated along the length of the villus has remained unresolved (*6*, *45*). Prevailing models have emphasized either mitotic pressure from crypt proliferation or local, cell-scale actomyosin dynamics as drivers of epithelial flow and turnover (*12*, *46*, *47*). Here, we identify a distinct organizing principle: Marangoni-like tissue flows driven by a geometry-encoded surface tension gradient and constrained by extracellular friction. By independently manipulating the driving surface tension gradient and extracellular friction, we show that epithelial velocity obeys the defining force balance of Marangoni flow, in which motion is set by the ratio of a tissue-scale tension gradient to resistive dissipation rather than by absolute force magnitude or local migratory activity. This framework explains how collective epithelial motion, spatially patterned extrusion, and tissue renewal emerge from the physical architecture of the villus.

Our findings resolve a long-standing paradox raised by prior studies. Live imaging experiments demonstrated that villus epithelial cells migrate actively rather than passively (*14*), yet did not identify the force field that organizes this movement over hundreds of microns. More recent organoid-based studies revealed that extrusion is regulated by contractility and mechanical competition, but lack native villus geometry and therefore cannot explain how extrusion is spatially patterned along the crypt-villus axis in vivo (*12*, *47*). By directly measuring nuclear strain in intact villi and linking it to curvature, adhesion and contractility, we show that tissue architecture itself generates a long-range surface tension gradient. This gradient gives rise to Marangoni-like flows that integrate migration and extrusion into a single tissue-scale physical mechanism.

A key insight from this work is that epithelial extrusion is most tightly coupled to the tension gradient itself, consistent with gradient-driven Marangoni flows in which material loss occurs at regions of maximal stress divergence rather than maximal stress (*48*). While crowding-driven extrusion operates in flat epithelia and simplified culture systems (*10*, *24*, *49*, *50*), our data show that in the context of the curved villus architecture, extrusion is a mechanically patterned process governed by tissue-scale force gradients. In Marangoni-like systems, gradients in surface tension naturally couple material flow to regions of release or dissipation. Similarly, in the villus epithelium, the tension gradient provides a mechanism by which continuous cell loss at the tip is synchronized with long-range collective motion from below (Table 3).

Critically, we further show that tissue-scale flows depend on a precise balance between driving and resisting forces (Table 3). While the surface tension gradient propels epithelial cells upward, adhesion to the ECM generates friction that opposes movement and must remain permissive for extrusion to occur. Increasing integrin-mediated adhesion elevates friction, collapses the effective tension gradient, and suppresses extrusion even when tissue tension is high (Table 3). Conversely, increasing contractility under high-friction conditions accelerates migration without restoring extrusion, leading to pathological cell accumulation (Table 3). These findings identify friction as a fundamental control parameter that determines whether Marangoni-like flows can sustain epithelial renewal.

The requirement for both a surface tension gradient and permissive friction reframes intestinal homeostasis as an emergent mechanical state rather than the outcome of any single cellular behavior (Table 3). Migration, extrusion, and tissue repair do not operate independently, but instead arise from a geometrically encoded force landscape that cells collectively experience and respond to. Disruption of either the gradient or the balance between tension and friction destabilizes this landscape and compromises renewal.

The physical picture emerging from our data is related to, but distinct from, previously described wetting and dewetting phenomena in epithelial cells (*48*, *51–53*). Prior work has shown that epithelial monolayers and aggregates can undergo active wetting or dewetting transitions driven by competition between traction, contractility, and adhesion, resulting in spreading or retraction events (*51*, *53*). These studies established that epithelia behave as active materials, and that global force balance governs large-scale morphological transitions. However, existing wetting frameworks primarily describe state transitions, whether a tissue spreads or retracts, and do not address how persistent, directional tissue flows can be maintained over long distances in a continuously renewing epithelium. In contrast, our work identifies a pre-existing, geometry-encoded surface tension gradient that persists under homeostatic conditions and continuously drives collective migration and extrusion along the villus axis. Rather than describing a transition between wet and dewet states, we show that intestinal renewal operates in a regime of gradient-driven, Marangoni-like flow, in which tissue architecture itself encodes the mechanical field that biases motion. This distinction is critical: whereas wetting describes whether a tissue spreads or retracts, the villus epithelium exploits geometry to sustain a stable, long-range flow that integrates migration and cell loss during homeostasis.

These findings suggest a new framework for understanding diseases characterized by villus blunting or architectural distortion, including inflammatory bowel disease and celiac disease (*54–56*). Disruption of villus architecture would therefore be expected to collapse the surface tension gradient, shift the balance between driving and dissipation, and suppress Marangoni-like epithelial flow even in the presence of intact cellular force generation. Defective epithelial renewal in these conditions may therefore reflect not only altered signaling or stem cell dysfunction, but also a failure to establish the mechanical gradients required for coordinated tissue-scale flow. For example, loss of villus curvature would be expected to collapse the surface tension gradient, disrupt epithelial flows, and uncouple migration from extrusion. Therapeutic strategies that restore tissue geometry or modulate contractility, adhesion, or ECM friction may help reestablish effective renewal.

More broadly, our results demonstrate how tissue geometry can encode long-range mechanical information that organizes collective cell behavior across space and time. By revealing Marangoni-like flows as a central mechanism of epithelial homeostasis, this work shifts the conceptual focus from local cytoskeletal activity to tissue-scale physical organization. Such geometry-driven mechanical flows may represent a general strategy by which rapidly renewing tissues maintain structure, function and homeostasis under constant turnover.

**Table 3.** Mechanical perturbations reveal a Marangoni-like force balance governing epithelial flow.

| **Perturbation** | **Primary mechanical effect** | **Expected effect on surface tension gradient ($\nabla\gamma$)** | **Expected effect on friction ($\xi$)** | **Observed effect on epithelial velocity** | **Interpretation** |
| --- | --- | --- | --- | --- | --- |
| Control | Native villus geometry and force balance | Preserved | Baseline | Stable, graded flow toward villus tip | Physiological Marangoni-like flow |
| DECMA-1 (E-cadherin inhibition) | Reduced cell-cell adhesion | Decreased | Approximately unchanged | Flow abolished | Loss of driving gradient collapses flow |
| Blebbistatin (myosin II inhibition) | Reduced actomyosin contractility | Decreased | Approximately unchanged | Flow abolished; tissue relaxation | Contractility required to sustain surface tension gradient |
| CK666 (Arp2/3 inhibition) | Disrupted actin-based force organization | Decreased | Approximately unchanged | Flow abolished | Active force organization required for gradient formation |
| 4-HAP (myosin II activation) | Enhanced actomyosin contractility | Increased | Approximately unchanged | Velocity increased | Flow scales with gradient magnitude |
| Mn<sup>2+</sup> (6 μM; moderate) | Increased integrin–ECM adhesion | Preserved | Increased | Velocity reduced | Increased dissipation suppresses flow |
| Mn<sup>2+</sup> (20 μM; high) | Strong integrin–ECM adhesion | Collapsed effective gradient | Strongly increased | Flow abolished | Dissipation-dominated regime |
| 4-HAP + Mn<sup>2+</sup> (6 μM) | Increased contractility with moderate friction | Increased | Increased | Partial rescue of velocity | Enhanced drive compensates moderate dissipation |
| 4-HAP + Mn<sup>2+</sup> (20 μM) | Increased contractility with high friction | Increased | Strongly increased | No rescue | Dissipation overwhelms driving force |
| Local blebbistatin photo-inactivation | Spatially reintroduced contractility | Local gradient imposed | Approximately unchanged | Flow reoriented toward high-tension region | Surface tension gradient sufficient to direct motion |

In Marangoni systems, material velocity is set by a balance between a surface tension gradient and resistive dissipation. Perturbations that reduce the tissue-scale tension gradient abolish epithelial flow, whereas increasing extracellular friction suppresses velocity despite preserved tension. Enhancing contractility compensates for increased friction only within a limited regime, demonstrating a Marangoni-like force balance governing epithelial motion.

## Acknowledgments

We thank members of the Sumigray lab for their insightful discussions; Valentina Greco and lab members for valuable scientific feedback and reagents.

## Funding

National Institutes of Health grant R35GM150645 (KS)

National Institutes of Health grant R01GR130179 (MPM)

American Cancer Society Research Scholar Grant (KS)

Sloan Matter-to-Life grant G-2025-79182 (MPM)

Human Frontiers Science Program grant RGP012/2025 (MPM)

Yale Stem Cell Center Chen Innovation Award (KS)

Yale Cancer Center Pilot Award (KS)

## Author contributions

Conceptualization: ZW, MPM, KS

Methodology: ZW, MPM, KS

Investigation: ZW

Visualization: ZW

Funding acquisition: KS, MPM

Project administration: KS

Supervision: MPM, KS

Writing – original draft: ZW, KS

Writing – review & editing: ZW, MPM, KS

## Competing interests

Authors declare that they have no competing interests.

## Data, code, and materials availability

The datasets, MATLAB code, and imaging data supporting the findings of this study are available from the corresponding author and will be deposited in a public repository upon publication.

## Supplementary Materials

Materials and Methods

Supplementary Text

Figs. S1 to S2

Tables S1

References (*1–60*)

Movies S1 to S17

## Materials and Methods

### Mice

All animal procedures were approved by the Yale University Institutional Animal Care and Use Committee and conducted in accordance with institutional guidelines. Mice were housed under a 12-hour light/dark cycle at a controlled ambient temperature of 22 ± 1 °C (72 °F ± 2 °F) and relative humidity of 50 ± 20%, with ad libitum access to food and water. Animals were maintained on a standard rodent diet (Inotiv Teklad Global 18% Protein Rodent Diet 2018S, autoclavable). Adult mice between 8 and 22 weeks of age were used for all experiments. Animals were euthanized prior to tissue collection according to approved protocols. The following mouse lines were used: mTmG (Jackson Laboratories, #007676)(*57*), ZO1-GFP(*58*) and Villin-rtTA (Jackson Laboratories, #031283)(*59*); mTmG; pTRE-H2B-GFP (Jackson Laboratories, #005104)(*60*). Both male and female mice were included unless otherwise specified.

### Live tissue culture medium

Basal medium was prepared using Advanced DMEM/F12 supplemented with 10 mM HEPES, 10% FBS, 1× ITS-G, and 1% penicillin–streptomycin. After thorough mixing, the medium was aliquoted into 50-mL tubes and stored at −20 °C for up to six months. On the day of use, one aliquot was thawed at room temperature and supplemented with 1× B27, 1× N2, and 10 mM N-acetylcysteine (NAC) to generate the complete medium. Complete medium was stored at 4 °C and used within seven days.

### Long-term live imaging

Jejunum was dissected from adult mice (8–22 weeks old) and immediately transferred into ice-cold Krebs buffer. After rinsing, tissues were moved to ice-cold Hibernate-A medium. Approximately 1-cm segments were embedded in low–gelling temperature agarose within a cryo-mold and placed at 4 °C for 5 min to solidify. Embedded samples were then mounted on a vibratome (tissue slicer) with the reservoir filled with ice-cold HBSS. Tissue slices were generated at 320 µm thickness using frequency and speed settings of 10. Slices were promptly transferred from the HBSS bath into Hibernate-A medium. For culture, 35-mm glass-bottom dishes were prepared with 300 µL of live tissue culture medium and a cell-insert placed in each dish. Tissue slices were positioned on the insert and incubated for 3 h at 37 °C with 5% CO₂. Prior to imaging, tissues were gently flushed with fresh live culture medium to remove dead cells, repositioned on the insert, and the dish medium was replaced with new medium containing 25 µM nifedipine. Dishes were then placed in a Tokai Hit GSXI2 environmental chamber (37 °C, 5% CO₂) mounted on a Leica Stellaris 5 confocal microscope equipped with a 25× water-immersion objective (0.95 NA). Samples were equilibrated for 30 min before imaging, and long-term live imaging was performed for 18 h.

### Drug treatment

Live tissue sections (prepared as described above) were treated under the following conditions: Control, DECMA-1 (15 µg/mL), blebbistatin (25 µM), CK666 (25 µM), and 4-hydroxyacetophenone (4HAP, 4 µM). Prior to drug application, tissues were gently flushed with live culture medium to remove dead cells, then incubated for 30 min in fresh medium containing nifedipine (25 µM). After this equilibration period, the medium was replaced with fresh nifedipine-containing medium supplemented with the respective drug (DECMA-1, blebbistatin, CK666, or 4-HAP). Tissues were imaged live for 18 h for velocity analysis. Following imaging, samples were fixed in 4% paraformaldehyde (PFA) for 20 min at room temperature for subsequent staining. To increase cell–ECM adhesion, manganese chloride (Mn²⁺) was applied at either 6 µM (low dose) or 20 µM (high dose). For manganese-treated live imaging, tissues were flushed and pre-incubated for 30 min in fresh medium containing nifedipine (25 µM), after which the medium was replaced with fresh nifedipine-containing medium supplemented with manganese. Dishes were equilibrated for 30 min in the Tokai Hit chamber before 18 h of live imaging. To assess the balance between tension and friction, combination treatments of manganese with 4HAP were performed. Tissues were first flushed to remove dead cells, then incubated for 30 min in medium containing manganese (6 µM or 20 µM). The medium was subsequently replaced with medium containing both manganese and 4HAP (4 µM), and tissues were immediately transferred to the microscope for 18 h of live imaging. Following imaging, samples were fixed in 4% paraformaldehyde (PFA) for 20 min at room temperature for subsequent staining.

### Tissue-scale laser ablation

Tissue-scale ablation was performed on an Andor Dragonfly 200 system equipped with a MicroPoint 405-nm laser and an Okolab stage-top incubator (5% CO₂, 37 °C). Tissue sections were prepared and maintained as described above. Ablation was carried out in control, DECMA-1, and 4HAP conditions. Adult mTmG mice were used. The ablation laser was operated in low-power mode at 90% intensity, 1 Hz, with 20 pulses per cycle for three cycles (total exposure: 60 s). Before ablation, z-stack live imaging was acquired to capture the 3D structure of the target villus. Following ablation, live imaging continued for 30 min to record tissue recoil and relaxation.

### Photo-inactivation of Blebbistatin

Blebbistatin photo-deactivation experiments were performed based on previous reports showing efficient inactivation with blue light, with 488 nm being the most effective wavelength (*39, 40*). Live tissues were first treated globally with blebbistatin to disrupt the endogenous tension gradient. Photo-deactivation was then carried out on a Leica Stellaris 5 microscope using the FRAP module. A 25× water-immersion objective with 2× digital zoom was used. A Z-stack of a single villus was acquired prior to stimulation. The FRAP cycle consisted of an imaging acquisition followed by illumination with the 488 nm laser at 25% intensity for eight pulses, after which imaging was resumed. This imaging–illumination cycle was repeated continuously for 30 minutes.

### Strain and strain gradient analysis

Nuclear strain measurement: Following 18 h of live imaging, tissues were fixed and stained as described above. Due to the strong inhibitory effect of blebbistatin, tissues in this condition were cultured for 3 h prior to fixation. After imaging, nuclei were segmented manually, as dense packing and z-projection made automated segmentation unreliable. Nuclear strain was defined as aspect ratio *(AR) − 1*. The position of each nucleus was mapped by measuring its linear distance from the villus tip along the epithelial axis. Nuclear strain values were plotted as a function of this distance for each villus.

Strain gradient analysis: Strain gradients were computed using the mapped nuclear strain profiles. The strain gradient between adjacent nuclei was defined as:

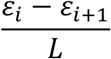

Where *ε*_i_ and *ε*_i+1_ are the strain values of neighboring nuclei and L is the inter-nuclear distance. Nuclear strain and position data were processed in custom MATLAB scripts to compute local strain gradients at 10 µm intervals along each villus. Strain-gradient profiles were averaged per villus to quantify intervillus variation. See supporting information for the code.

### Tissue elongation analysis

To quantify tissue elongation following myosin inhibition, epithelial cell movements along the villus axis were analyzed immediately after blebbistatin application. Individual cells were tracked over time, and their displacement along the villus axis was computed from time-lapse imaging. Elongation was quantified over the first 3 hours following drug addition, a time window that captures the full elongation response, as villus extension reached a stable value within this period and did not increase further at later times. Tissue elongation was expressed as a dimensionless strain by normalizing cell displacement to the initial villus length *L_0_*, measured prior to drug treatment. For each cell, cumulative displacement *L*_(t)_was calculated as a function of time and normalized according to

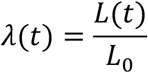

This normalized displacement provides a spatially resolved proxy for tissue stretch induced by removal of myosin-generated contractile forces. For each villus region, strain values were averaged across all tracked cells to obtain the mean regional elongation as a function of time. The total elongation for each region was defined as the change in mean strain between the initial state and the fully elongated state reached within the 3-hour window. This elongation amplitude reflects the extent of mechanical extension following myosin inhibition and therefore serves as a readout of pre-existing contractile stress within the tissue. Villus tip regions consistently exhibited greater elongation than basal regions, indicating higher stored contractile energy localized to the villus tip.

### Immunofluorescent staining

Live tissues were fixed in 4% PFA for 20 mins at room temperature. Next, PFA was removed, and tissues were washed by PBS for three times, 5 minutes each time, on the rocker. And then the tissues were blocked by 1% BSA, BSA was dissolved in PBST (PBS with 0.2% Tween), for 2 hours at room temperature. Then BSA with PBST was removed. Tissues were incubated with primary antibody in BSA with donkey serum and was incubated in 37 °C for two days. Next, the primary antibody was washed using PBST on a rocker, for 3 times, 5 min for each time. Following secondary antibody. The secondary antibody was also in PBST with BSA and donkey serum, then incubate in 37 °C for one day. Next, PBST was used to wash the tissue. Three times wash and 5 minutes each time. Then tissue was taken to a Leica Stellaris 5 to take imaging.

### Image registration

Time-lapse image stacks were corrected for global tissue drift prior to analysis. Raw TIFF image sequences were converted to double precision and aligned frame-by-frame using a rigid translation model. For each frame, an average reference image constructed from preceding corrected frames was used to estimate translational offsets via normalized cross-correlation. Coarse alignment was subsequently refined using optical flow–based motion estimation to correct residual subpixel drift, with corrections constrained to small displacements to prevent overcorrection. Frames were resampled using cubic interpolation and saved as a drift-corrected image stack for downstream cell tracking and tissue-scale analysis. This procedure corrects global tissue drift while preserving relative cellular motion (see Code Availability).

### Cell tracking

Cell trajectories were obtained using the MTrackJ plugin in Fiji and exported as CSV files containing track identifiers (TID), frame indices (PID), and centroid coordinates (x,y). Trajectories were reconstructed by grouping coordinates by track ID and ordering positions by frame number. Cell displacements were expressed relative to each cell’s initial position. Instantaneous velocity vectors were computed from frame-to-frame centroid displacements divided by the imaging interval Δ*t*, and cell speed was calculated as the magnitude of the velocity vector. For each cell, mean migration speed was defined as the time-averaged speed, and total travel distance was computed as the cumulative sum of frame-to-frame displacements (see Code Availability).

### Data analysis and Statistics

Cell velocity analysis: Villus length was normalized from 0 to 1 (tip to base) and subdivided into five equal regions (0–0.2, 0.2–0.4, 0.4–0.6, 0.6–0.8, 0.8–1.0). For each region, a 100 × 100 µm field of view was extracted. Cells were tracked over 3 h of live imaging. Cell motion was semi-manually tracked using MTrackJ (Fiji). The software output x–y coordinates for each cell at every frame over the 3 h period. Coordinate data were processed in custom MATLAB scripts to calculate mean cell velocity for each cell. The mean velocity for each villus was plotted as a function of normalized position along the villus (0.2, 0.4, 0.6, 0.8, 1.0). For each condition, velocities were averaged per villus to capture intervillus variability.

## Antibodies

Primary antibodies: anti-E-Cad (13-1900, Invitrogen, 1:100), anti-Myosin IIA (909802, BioLegend, 1:100).

Secondary antibodies: Alexa Fluor 647 donkey anti-rat IgG antibody (Invitrogen, 1:500), Alexa Fluor 488 donkey anti-rabbit IgG antibody (Invitrogen, 1:500).

## Notes

### Competing Interest Statement

The authors have declared no competing interest.

