## Supplemental Materials for "Tissue geometry encodes a surface tension gradient that drives epithelial renewal"

**The PDF file includes:**

Figs. S1 to S2
Captions for Movies S1 to S17

**Other Supplementary Materials for this manuscript include the following:**

Movies S1 to S17 (.mp4)

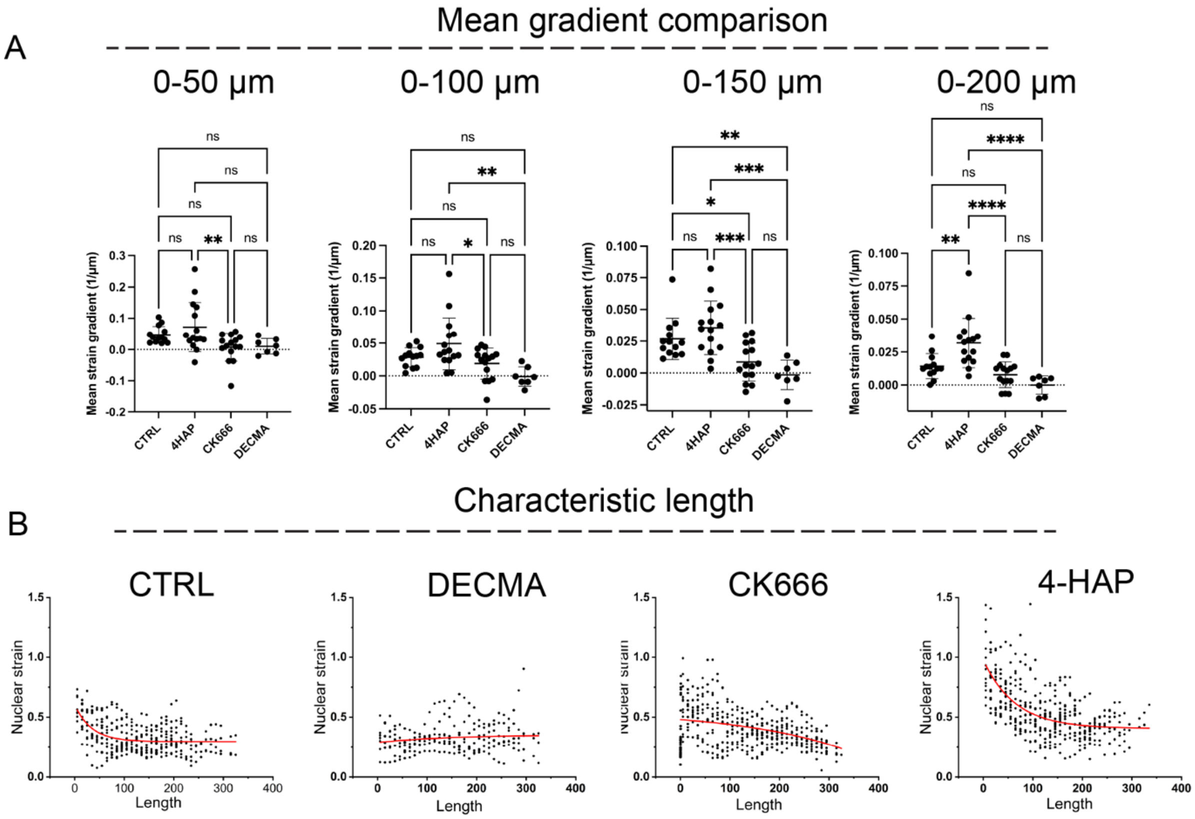


**Fig. S1. Large-scale tension gradient exists in the villus axis.** A) Mean nuclear strain gradient quantified over increasing distance ranges from the villus tip (0–50 µm, 0–100 µm, 0–150 µm, and 0–200 µm) across all experimental conditions. Statistically significant differences between conditions first emerge when gradients are averaged over ≥150 µm from the villus tip. B) Characteristic decay length of nuclear strain along the villus axis estimated by fitting an exponential decay model ($\boldsymbol{\varepsilon} = \boldsymbol{a} * \boldsymbol{e}^{-\frac{l}{L}} + \boldsymbol{\varepsilon_0}$), where $\boldsymbol{\varepsilon}$ is nuclear strain at position $l$ along the villus axis measured from the tip, $\boldsymbol{\varepsilon_0}$ is the strain at the villus tip, $a$ is a fitted constant, and $L$ represents the characteristic length scale of the tension-dominated region. Data are mean ± SD. Multigroup comparisons used one-way analysis of variance with a Tukey test.

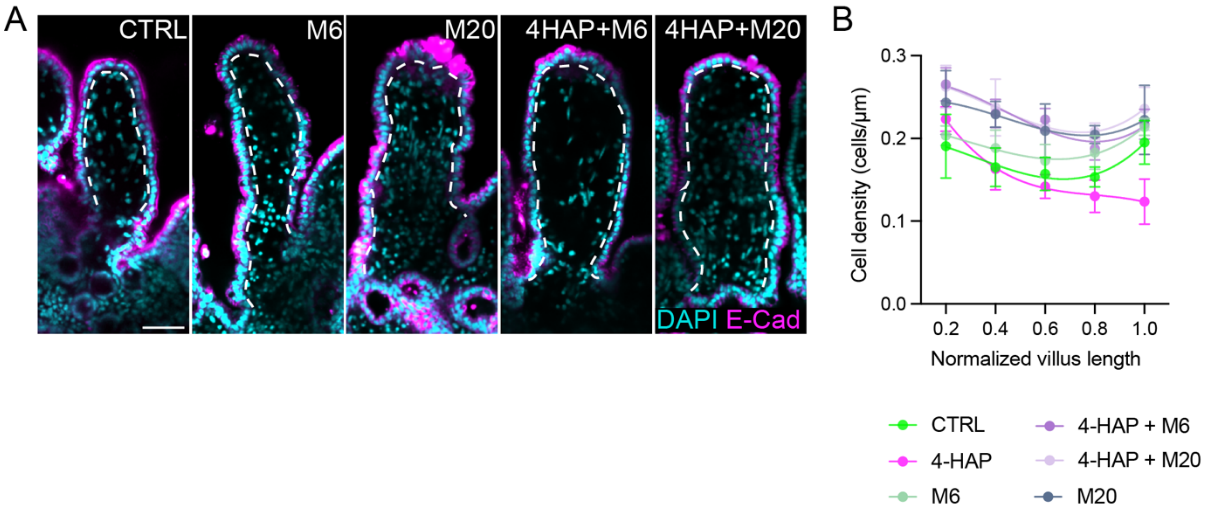


**Fig. S2. Tissue structure and density upon perturbed balance between surface tension gradient and extracellular friction. A)** Representative confocal images of tissue treated 24 hours with the indicated conditions. Scale bar, 50µm. **B)** Cell density measurement along villus axis. Data are mean ± SD.

Movie S1.
Long-term live imaging of ex vivo intestinal tissue sections showing preservation of villus architecture and sustained physiological epithelial turnover during extended culture. Time is shown as hours:minutes (h:min).

**Movie S2.**
High-magnification live imaging showing the dynamics of epithelial cell extrusion at the villus tip during homeostatic renewal. Time is shown as hours:minutes (h:min).

**Movie S3.**
High-magnification live imaging showing the dynamics of epithelial cell mitosis and reintegration in the crypt during homeostatic renewal. Time is shown as hours:minutes (h:min).

**Movie S4.**
Live imaging showing intestinal epithelial cells exhibit coherent directional flow along the villus axis. Time is shown as hours:minutes (h:min).

**Movie S5.**
Live imaging of DECMA-1–treated intestinal tissue showing loss of coordinated epithelial flow due to impaired cell–cell adhesion. Time is shown as hours:minutes (h:min).

**Movie S6.**
Live imaging of Blebbistatin–treated intestinal tissue showing loss of coordinated epithelial flow due to impaired actomyosin contractility. Time is shown as hours:minutes (h:min).

**Movie S7.**
Live imaging of CK666–treated intestinal tissue showing loss of coordinated epithelial flow due to impaired Arp2/3-dependent actin assembly. Time is shown as hours:minutes (h:min).

**Movie S8.**
Live imaging of 4-HAP–treated intestinal tissue showing enhanced coordinated epithelial flow driven by increased actomyosin contractility. Time is shown as hours:minutes (h:min).

**Movie S9.**
Live imaging of intestinal tissue treated with blebbistatin showing rapid tissue relaxation and elongation during the first 3 hours of imaging. Time is shown as hours:minutes (h:min).

**Movie S10.**

Live imaging of intestinal tissue treated with manganese (6 μM), showing reduced coordination of epithelial flow associated with increased extracellular matrix–derived friction. Time is shown as hours:minutes (h:min).

**Movie S11.**

Live imaging of intestinal tissue treated with manganese (20 μM), showing significantly reduced coordinated epithelial flow associated with increased extracellular matrix–derived friction. Time is displayed as hours:minutes (h:min).

**Movie S12.**

Live imaging of intestinal tissue treated with 4-HAP and manganese (6 μM), showing partial restoration of coordinated epithelial flow. Time is displayed as hours:minutes (h:min).

**Movie S13.**

Live imaging of intestinal tissue treated with 4-HAP and manganese (20 μM), showing failed restoration of coordinated epithelial flow. Time is displayed as hours:minutes (h:min).

**Movie S14.**

Representative long-term live imaging of intestinal tissue treated with 4-HAP, showing cell extrusion at the villus tip over 14 hours. Cyan, cell nuclei; magenta, cell membranes. Time is displayed as hours:minutes (h:min).

**Movie S15.**

Live imaging of intestinal tissue following laser ablation, illustrating wound opening and closure dynamics. Time is displayed as minutes:seconds (min:s).

**Movie S16.**

Live imaging of intestinal tissue treated with 4-HAP following laser ablation, illustrating enhanced wound opening and closure dynamics. Time is displayed as minutes:seconds (min:s).

**Movie S17.**

Live imaging of intestinal tissue treated with DECMA-1 following laser ablation, illustrating wound opening and closure dynamics. Time is displayed as minutes:seconds (min:s).
